# HIRA defines early replication initiation zones independently of their genome compartment

**DOI:** 10.1101/2024.08.29.610220

**Authors:** T. Karagyozova, A. Gatto, A. Forest, J.-P. Quivy, M. Marti-Renom, L. Mirny, G. Almouzni

## Abstract

Chromatin states and 3D architecture have been used as proxy to identify replication initiation zones (IZs) in mammalian cells. While they do often correlate, their functional interconnections remain a puzzle. Here, we dissect these relationships by focusing on the histone H3.3 chaperone HIRA, which plays a role in both early initiation zone (IZ) definition and higher-order organisation of active chromatin. We monitored in parallel early replication initiation, chromatin accessibility, histone post-translational modifications (PTMs) and 3D organisation in wild-type cells, HIRA knock-out cells and HIRA knock-out cells complemented with HIRA. In the absence of HIRA, impaired early firing at HIRA-dependent IZs does not correspond to changes in chromatin accessibility or patterns of histone H3 PTMs. With respect to 3D organisation, a small subset of early IZs initially in compartment A switched to B and lost early initiation in the absence of HIRA. Critically, HIRA complementation restores these early IZ (and H3.3 variant enrichment) without substantial compartment reversal. Thus, our work reveals that regulation of early replication initiation by HIRA can be uncoupled from accessibility, histone mark patterns and compartment organisation.

## Introduction

The synthesis of new DNA starts at the origins of replication: estimated 30,000-50,000 sites in the mammalian genome, licensed in G1 before only a small subset are used stochastically in S phase (Méchali, 2010). In metazoans, origin definition combines properties of DNA sequence (Cayrou et al., 2011; Dellino et al., 2013) and epigenetic features (Cadoret et al., 2008; MacAlpine et al., 2010; Miotto et al., 2016). Initiation events are typically clustered in zones (IZs) of 20-150kb (Cadoret et al., 2008; Cayrou et al., 2011; Petryk et al., 2016; Wang et al., 2021; Zhu and Kanemaki, 2024), which do not fire at the same time, but follow a spatiotemporal order of activation, termed the replication timing (RT) programme (Vouzas and Gilbert, 2021). RT is cell-type specific and strongly correlates with chromatin state and its three-dimensional (3D) organization (Vouzas and Gilbert, 2021). Typically, early-replicating regions are transcribed (Dellino et al., 2013), accessible (Sugimoto et al., 2018), decorated by active histone marks (Cayrou et al., 2015) and correspond to compartment A identified by Hi-C (Hansen et al., 2010; Pope et al., 2014; Rivera-Mulia et al., 2015). Few exceptions to these correlations have been identified in early development (Dileep et al., 2019; Miura et al., 2019; Nakatani et al., 2023), but RT and chromatin features are not fully established at this stage, making it challenging to translate this in the context of differentiated cells. Thus, a major question in the field has been to disentangle the functional connections between early replication, epigenetic state, and 3D chromatin organization.

Recently, a role for histone variants has also emerged in the context of the regulation of replication initiation (Gatto et al., 2022; Karagyozova and Almouzni, 2024; Long et al., 2020). With respect to histone H3, we first found that enrichment pattern of H3.3 and H3.1 followed early versus late RT respectively (Clément et al., 2018). The replicative H3.1 variant, produced at high levels at S phase entry (Franklin and Zweidler, 1977), is deposited in a DNA synthesis-coupled (DSC) manner (Tagami et al., 2004). This is mediated by the CAF-1 complex (Smith and Stillman, 1989; Tagami et al., 2004), coupled to replisome progression through its interaction with the sliding clamp PCNA (Moggs et al., 2000). In contrast, the replacement variant H3.3 is expressed throughout the cell cycle (Wu and Bonner, 1981) and incorporated in a DNA synthesis-independent (DSI) manner (Drané et al., 2010; Tagami et al., 2004). The first identified chaperone involved in this DSI pathway is the HIRA complex (Ray-Gallet et al., 2002; Tagami et al., 2004). HIRA ensures H3.3 enrichment at active regions, regulatory elements (Goldberg et al., 2010; Ray-Gallet et al., 2011) and sites of high turnover (Deaton et al., 2016), likely through interacting with RNA Pol II (Ray-Gallet et al., 2011). Additionally, HIRA deposits H3.3 at transiently exposed DNA in a gap-filling manner (Ray-Gallet et al., 2011). More recently, by mapping *de novo* incorporation of H3.1 and H3.3 in S phase (Gatto et al., 2022), we revealed that new H3.3 deposition occurred systematically at pre-existing H3.3-enriched sites, while new H3.1 followed the replication fork movement. This dual deposition mechanism results in the establishment of H3.3/H3.1 boundaries. Strikingly, these boundaries overlap with early replication IZs. Notably, HIRA knock-out (KO) disrupted not only these H3.3/H3.1 boundaries, but also the corresponding replication IZs independently of transcription (Gatto et al., 2022), uncovering an unanticipated connection for HIRA and DNA replication. In the absence of HIRA, we distinguished two types of early IZs: (i) those that we called “blurred sites” since both the H3.3 pattern and replication initiation of the early IZs became fuzzy at the boundaries of H3.3 peaks and (ii) those that we called buried sites corresponding to singular H3.3 peaks which disappeared along with abrogation of replication initiation (Gatto et al., 2022). “Blurred sites” and “buried sites” corresponded respectively to actively transcribed and inactive domains. The next major issue was then to understand which functional relationships could link H3.3 deposition by HIRA and early replication. Recently, we uncovered an importance of HIRA in higher-order organisation of active chromatin (Karagyozova et al., 2024). We found that HIRA is key to maintain compartment A interactions and the local organisation of active genes by modulating chromatin accessibility (Karagyozova et al., 2024). Thus, histone H3.3 chaperone HIRA plays a role in both early initiation zone (IZ) definition and higher-order organisation of active chromatin. These findings prompted us to consider whether HIRA could actually contribute to the definition of early IZs by impacting their higher-order organisation.

Here, we combined knock-out (KO) and rescue experiments of the histone H3.3 chaperone HIRA to dissect the relationship between early replication initiation, chromatin state, and 3D genome organisation. First, we demonstrated that HIRA defines early IZs independently from its impact on accessibility and without affecting their H3 PTM pattern. Second, we found that HIRA-dependent non/low-transcribed early IZ (buried sites) do not exclusively belong to compartment A in wild-type (WT) cells. Furthermore, in the absence of HIRA, we also found that a fraction of IZs initially in compartment A switched to B as they lost capacity to initiate early replication. Yet, rescue with HIRA recovered the H3.3 enrichment and early firing patterns at early IZs without necessarily restoring their compartment A identity. Thus, our results indicate that HIRA defines early IZs independently of their accessibility, PTM pattern and compartment organisation.

## Results

### HIRA defines early replication IZs independently of their accessibility

To understand the functional interplay between early replication and H3 variant balance, we explored concomitantly chromatin state and higher-order organization. We focused on the HIRA-dependent early replication initiation zones (IZs) we previously identified and coined blurred and buried (Gatto et al., 2022) (Figure 1A for schematic illustration and Figure 1B for representative tracks of H3.3/H3.1 balance and early replication). Given the links between early replication (Cayrou et al., 2015), H3.3 (Martire et al., 2019) and open chromatin, and our recent work on the importance of HIRA in regulating accessibility in active chromatin domains (Karagyozova et al., 2024), we first examined if HIRA affected early IZ firing by modulating their accessibility. Using sites from (Gatto et al., 2022) as a reference (scheme in Figure 1A), we probed chromatin accessibility in WT and HIRA KO cells by ATAC-Seq and compared it to H3.3 and early replication patterns (Figure 1C). In WT cells, we detected high accessibility at the boundaries for both types of sites (corresponding to early IZs) in WT cells (Figure 1C, Supplementary Figure 1A). In the absence of HIRA, within blurred sites, ATAC-seq signal showed a striking increase of accessibility (Figure 1C). However, ATAC-seq signal maintained sharpness at their boundaries (Supplementary Figure 1A) and did not mirror the blurring of H3.3 enrichment (Figure 1C), H3.3/H3.1 ratio (Supplementary Figure 1B) and EdU 2h signal (Figure 1C). At buries sites, accessibility weakly decreased in the absence of HIRA, while maintaining its pattern (Figure 1C, Supplementary Figure 1A). This was in sharp contrast with the complete loss of H3.3 enrichment and early firing (Figure 1C) and was not matched by changes in expression (Supplementary Figure 1B). These data suggest that the impact of HIRA on early initiation is independent of DNA accessibility. To confirm this, we also analysed H3.3-rich early IZs identified by OK-seq at 1kb resolution (Gatto et al., 2022; Petryk et al., 2016). Indeed, both H3.3 and EdU 2h enrichment blurred and decreased at non/low-expressed H3.3^+^ OK-seq IZs without a corresponding change in ATAC-seq in the absence of HIRA (Supplementary Figure 1C). In contrast, accessibility increased at early IZs flanked by transcribed sites scaling with their expression level. Thus, we can conclude that HIRA regulates early firing at both blurred and buried sites independently of chromatin accessibility.

**Figure 1.**
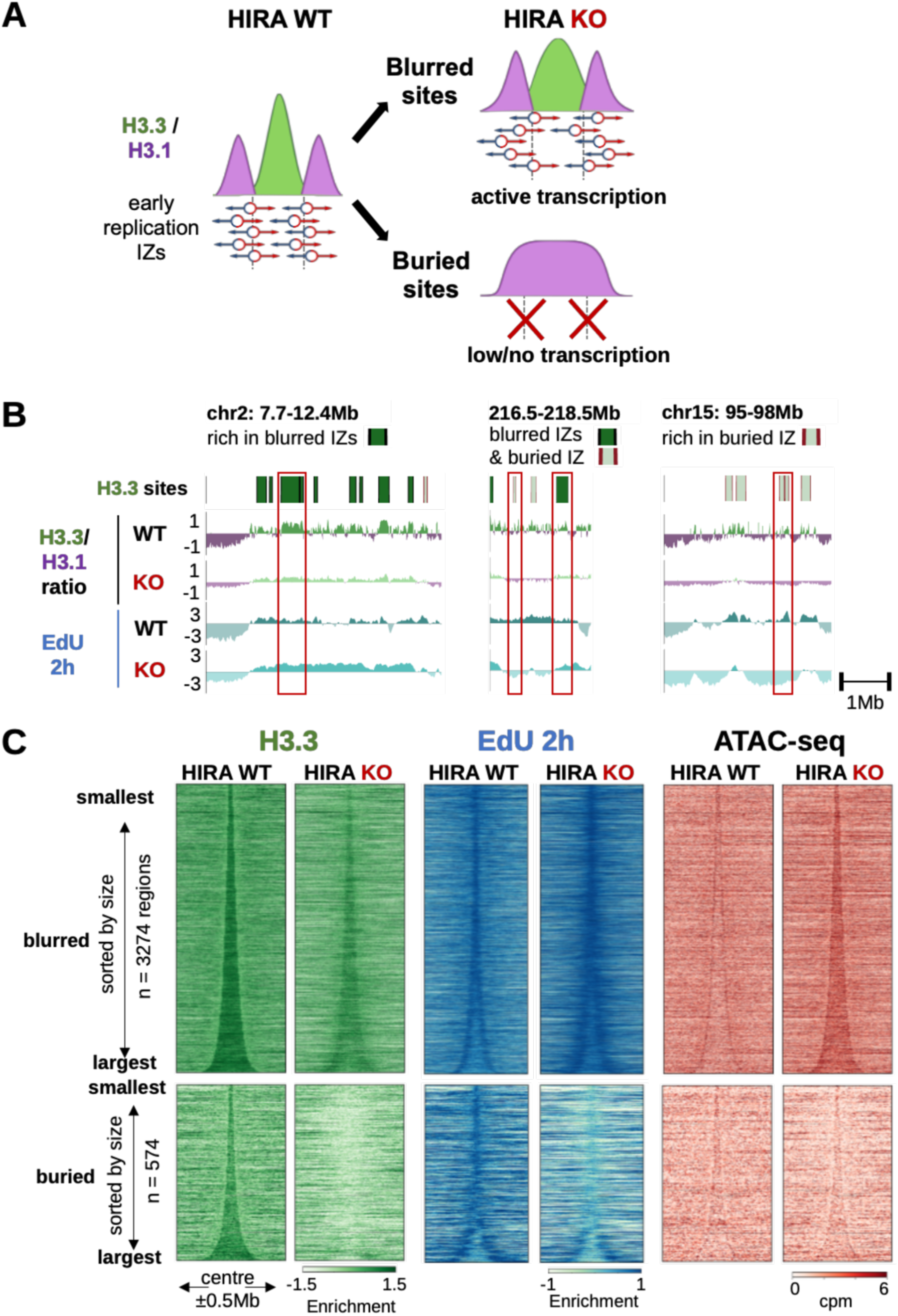
HIRA defines early replication initiation zones independently of its importance for chromatin accessibility. **A.** Schematic representation of early replication initiation zones defined by HIRA-dependent H3.3/H3.1 boundaries as described in (Gatto et al., 2022). **B.** H3.3/H3.1 ratio, and early replication (EdU 2h) from WT and HIRA KO cells at representative chromosomal regions containing blurred and buried sites (denoted above the tracks in dark and light green, respectively). Individual examples of blurred and buried sites are highlighted in red boxes in each representative region. **C.** Enrichment of H3.3, EdU at 2h in S and ATAC-seq signal from WT and HIRA KO cells at blurred (n = 3274, top) and buried sites (n = 574, bottom), sorted by size and centered at their middle ±0.5Mb. Enrichment of H3.3 and EdU is shown as z-score of log_2_ IP/input, ATAC-seq is shown as cpm at 10kb bins.

### Impaired early IZ firing in the absence of HIRA is not associated with local changes in H3 marks

Early IZs have been associated with the presence of both active and inactive H3 marks (Cayrou et al., 2015). Given its links with transcription, H3.3 itself often coincides with active marks (Goldberg et al., 2010). Furthermore, phosphorylation of the unique H3.3S31 in interphase can enhance H3K27ac at enhancers (Armache et al., 2020; Martire et al., 2019; Morozov et al., 2023). In light of these connections, we examined the effect of HIRA loss at early IZs on a set of selected active (H3K4me3, promoter-associated, H3K4me1, H3K27ac, enhancer-associated) and inactive (H3K9me3, constitutive heterochromatin and H3K27me3, facultative heterochromatin) PTMs profiled by native ChIP-seq (Karagyozova et al., 2024). In WT cells, H3K4me1 and H3K27ac were enriched, while H3K9me3 and H3K27me3 were depleted at both types of sites (Figure 2A). The active promoter mark H3K4me3 was enriched at the boundaries of blurred, but not buried sites, in line with their link with transcription (Gatto et al., 2022). In the absence of HIRA, we did not detect reduced precision or drastic redistribution of the marks matching the changes in H3.3 and early firing at either type of site (Figure 2A, B). In terms of enrichment, we detected small changes predominantly at buried sites, but they were not comparable to the extent of H3.3 loss in the absence of HIRA (Supplementary Figure 2A). To confirm these results, we also analysed PTM patterns at H3.3^+^ OK-seq IZs at 1kb bins (Supplementary Figure 2B) and again we did not observe changes that compared to H3.3 or EdU 2h (Supplementary Figure 1C). Thus, we can conclude that the importance of HIRA in defining early replication initiation zones does not rely on histone PTMs.

**Figure 2.**
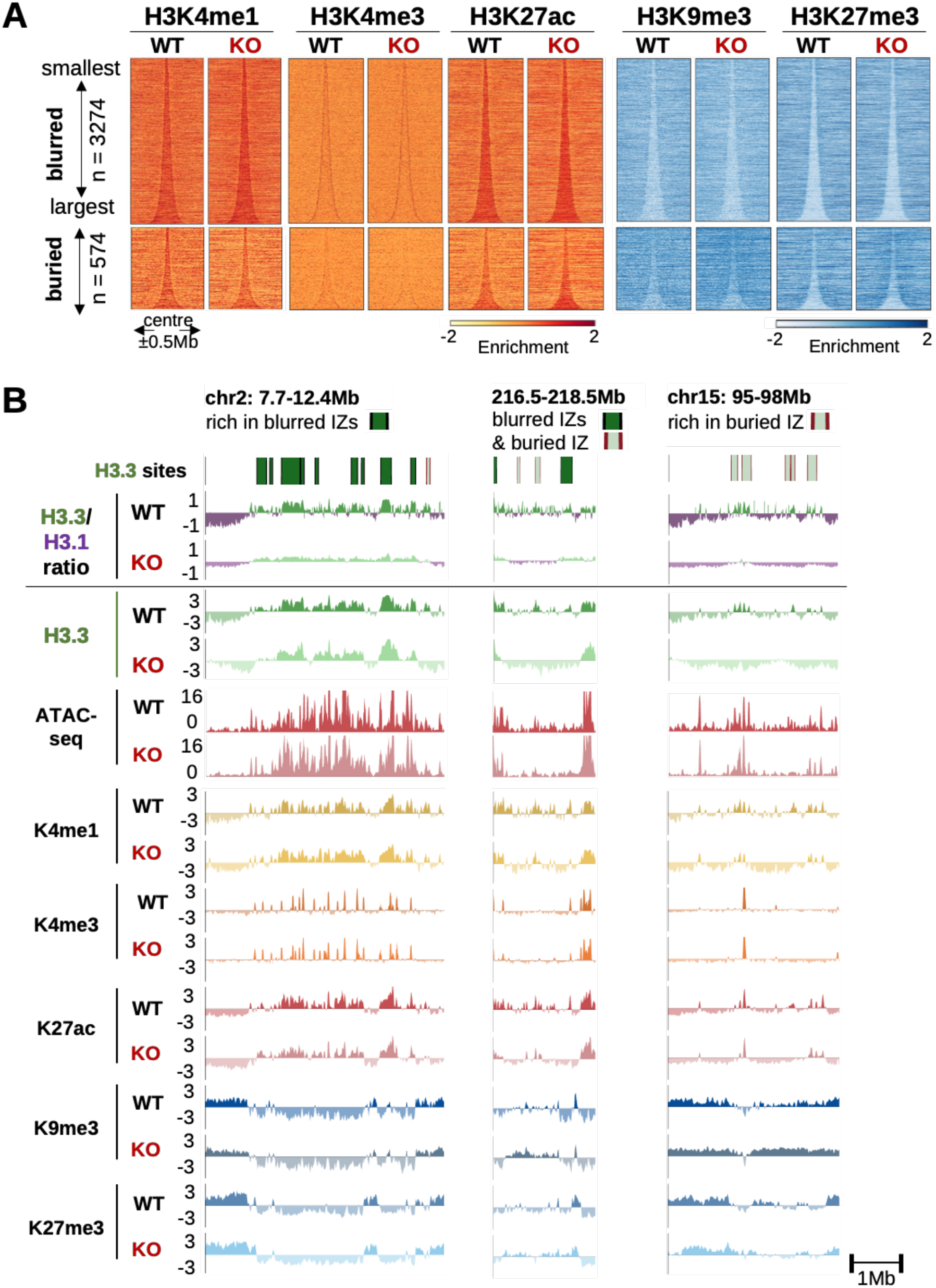
Impaired early IZ firing in the absence of HIRA is not associated with local H3 PTM redistribution. **A.** Active (H3K4me1, H3K4me3, H3K27ac) and repressive (H3K9me3, H3K27me3) histone PTM enrichment profiles from WT and HIRA KO cells at blurred (n = 3274, top) and buried (n = 574, bottom) sites, centered at their middle ±0.5Mb and sorted by size. **B.** H3.3/H3.1 ratio (for reference, as in Figure 1B), H3.3 enrichment, ATAC-seq signal and enrichment of active (H3K4me1, H3K4me3, H3K27ac) and repressive (H3K9me3, H3K27me3) histone PTMs from WT and HIRA KO cells at representative chromosomal regions (as in Figure 1B) containing blurred and buried sites (denoted above the tracks in dark and light green, respectively). Enrichment shown is z-score of log_2_ IP/input, ATAC-seq signal is shown as cpm at 10kb bins. In the representative tracks, signals are shown at 10kb bins smoothed over 3 non-zero bins.

### In the absence of HIRA, only non-transcribed early IZs switch from compartment A to B

In addition to chromatin state, early firing also correlates with features of higher-order genome organisation. More specifically, early replicating regions generally correspond to compartment A, and an enrichment of early IZs has been reported at TAD borders (Vouzas and Gilbert, 2021). Using Hi-C, we recently observed that HIRA promotes maintenance of compartment A interactions without affecting TAD organisation. Furthermore, HIRA loss leads to minor compartment switching (3.1% genome) independently of transcription (Karagyozova et al., 2024). Thus, we wondered whether HIRA can impact firing at early IZs by influencing their spatial arrangement. We examined the compartment identity of blurred and buried sites based on the first eigenvector (EV1) from Hi-C maps, where positive values correspond to compartment A and negative, to B (Lieberman-Aiden et al., 2009). Blurred sites were generally in compartment A in both WT (85% sites) and HIRA KO cells (86% sites, Figure 3A, green points). This was also reflected by their positive EV1 values in both conditions (Figure 3B) and absence of substantial compartment switching compared to random sites (Figure 3C). In contrast, buried sites were mainly in compartment B in WT (67% sites, Figure 3A, red points, B). Furthermore, buried sites significantly switched compartment in HIRA KO (23.5%, Figure 3C, Supplementary Figure 3A), predominantly from A to B (20%, Figure 3D). Given the association of compartment A with gene expression, we examined transcription at buried sites by RNA-seq to understand if it could explain their behaviour. In WT cells, buried sites were in either compartment A or B regardless of their expression (Figure 3D). Buried sites which switched from A to B had significantly lower expression in WT cells than the ones that remain in compartment A in the absence of HIRA (Figure 3D, Supplementary Figure 3B). However, A-to-B switching in HIRA KO was not accompanied by a significant decrease in transcription (Supplementary Figure 3B, C). In terms of PTMs, H3K9me3 and H3K27me3 increased while H3K4me1 decreased at A-to-B buried sites in HIRA KO cells (Supplementary Figure 3D). However, these changes were weaker than the depletion of H3.3 (Supplementary Figure 3D). Thus, our data reveal that transcribed early IZs (blurred sites) are found in compartment A irrespective of HIRA status. In contrast, low/non-transcribed early IZs (buried sites) are predominantly in compartment B in WT cells, indicating that early initiation does not always occur in compartment A in differentiated cells. Furthermore, only non-transcribed buried sites switched from compartment A to B along with loss of early firing in the absence of HIRA.

**Figure 3.**
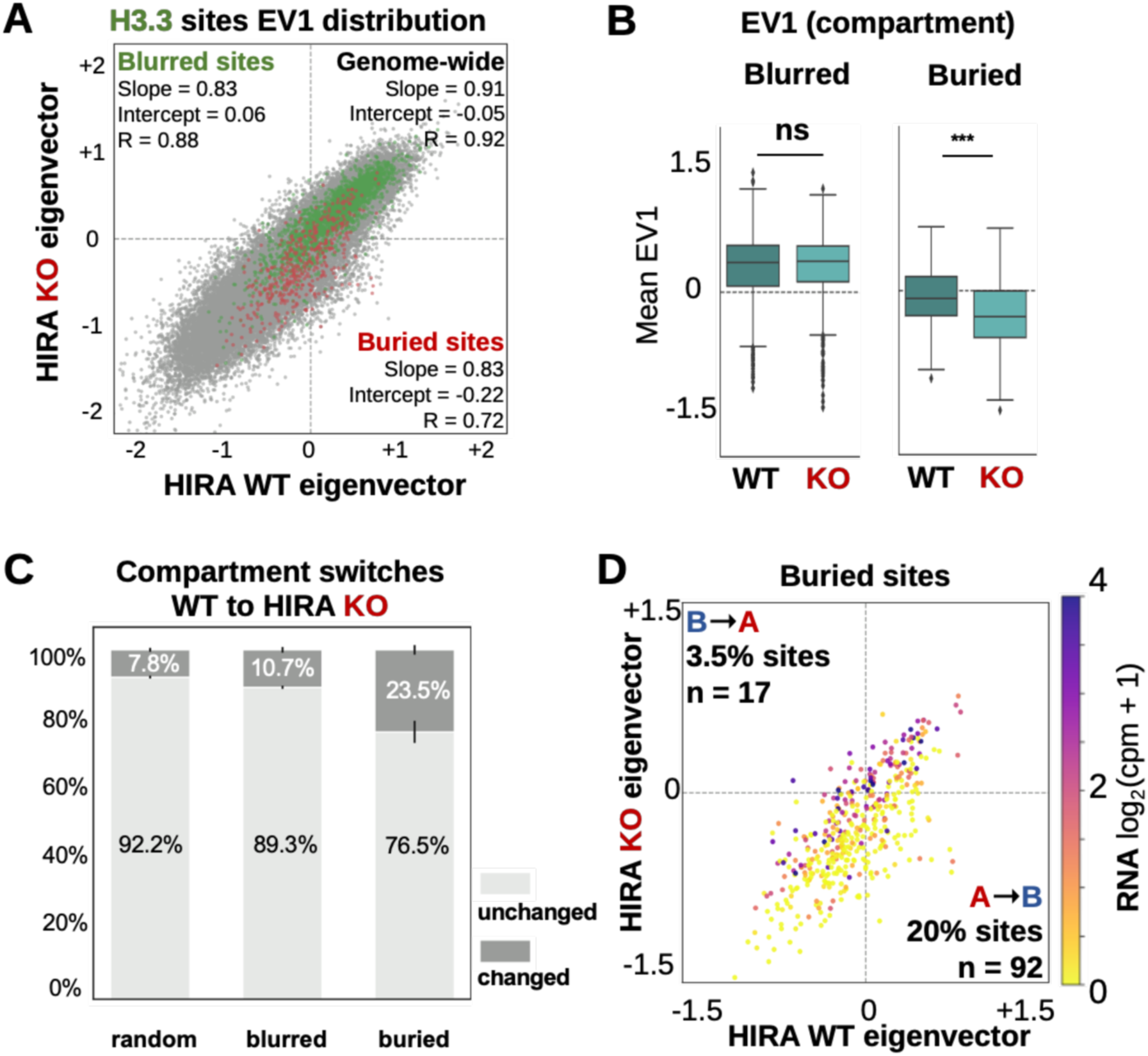
In the absence of HIRA, only non-transcribed early IZs switch from compartment A to B. **A.** Comparison of EV1 distribution genome-wide (50kb bins, n = 40887, grey) or at blurred (n = 2382, green) and buried (n = 439, red) sites between WT and HIRA KO cells. Slope, intercept and R from linear regression are noted for each set. **B.** Mean value of EV1 (1^st^ eigenvector, indicating compartment) at blurred and buried sites from WT and HIRA KO cells. **C.** Proportions of blurred and buried sites which remain in the same compartment (unchanged) or undergo a switch (changed) from WT to HIRA KO cells. A set of randomised size-matched sites was quantified as control. **D.** Scatterplot of mean EV1 value at buried sites (n = 439) from WT and HIRA KO cells. Colour represents transcriptional activity from HIRA WT cells. Proportion of sites which change from compartment A-to-B (lower right quadrant) or B-to-A (upper left quadrant) are quantified. EV1 was calculated from 50kb-binned Hi-C matrices and then re-binned to 10kb to compute mean EV1 value for blurred and buried sites. Transcriptional activity is measured as the mean log_2_(cpm+1) RNA-seq signal binned at 10kb for each site. Two-tailed Mann-Whitney U test corrected for multiple testing by FDR (5% cut-off) was used to determine significance of differences between WT and HIRA KO. Significance was noted as: * (p<=0.05), ** (p<=0.01), *** (p<=0.001) for all comparisons.

### HIRA rescue reestablishes H3.3 pattern and early replication initiation at both blurred and buried sites

To determine if reestablishing precise H3.3 deposition can restore both timely firing at early replication sites and compartment organization, we performed rescue experiments. We transiently transfected HIRA-YFP (HIRA) or YFP only (control) plasmid for 48h in H3.1-SNAP and H3.3-SNAP HIRA KO cells followed by G1/S-synchronization (Figure 4A), obtaining high efficiency (>70%) in all conditions (Supplementary Figure 4A). We then evaluated H3.3 and H3.1 distribution at blurred and buried sites by SNAP-Capture ChIP-seq. Rescue with HIRA, but not with control plasmid, restored both the pattern and levels of H3.3 enrichment at blurred sites (Figure 4B, C, top, Supplementary Figure 4B). Strikingly, HIRA complementation was also sufficient to target H3.3 incorporation to buried sites, despite their complete loss of H3.3 upon HIRA KO and low or absent transcriptional activity (Figure 4B, C, bottom, Supplementary Figure 4B). Concomitantly, we also detected recovery of the H3.1 enrichment and the H3.3/H3.1 ratio (Supplementary Figure 4C, D). Thus, re-supplying HIRA is sufficient to reestablish H3.3 enrichment and H3.3/H3.1 balance at blurred and buried sites.

**Figure 4.**
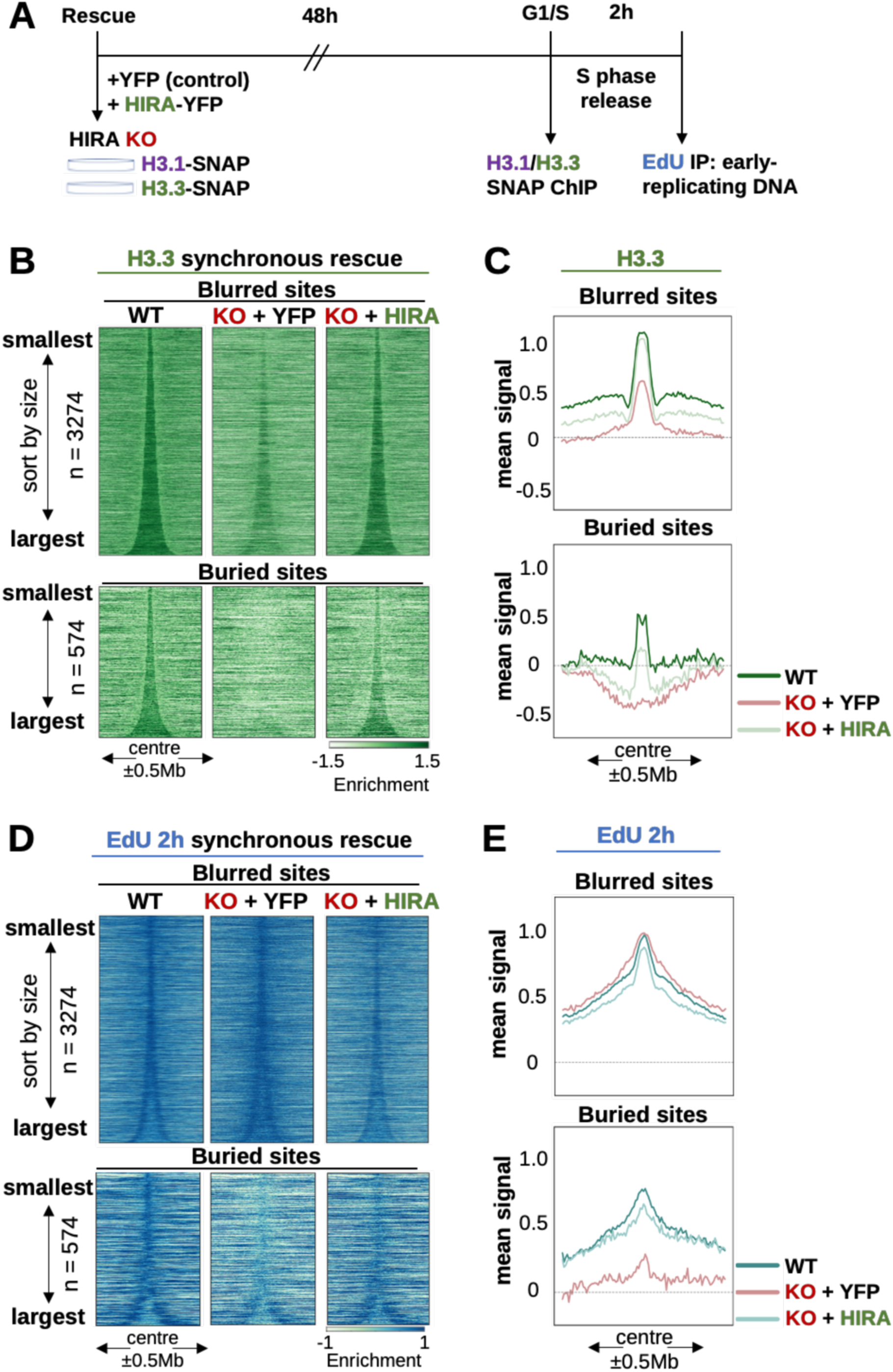
HIRA rescue reestablishes H3.3 pattern and early replication initiation at both blurred and buried sites. **A.** Scheme of experimental strategy to perform HIRA rescue combined with G1/S synchronization to assay total H3.1/H3.3-SNAP by ChIP-seq and new DNA synthesis in early S phase (2h release). Asynchronous cells constitutively expressing H3.1- or H3.3-SNAP were transfected with YFP (control) or HIRA-YFP plasmid. Cells were then arrested at the G1/S boundary by double thymidine block (starting 6h post-transfection). Total H3.1- and H3.3-SNAP were assayed by SNAP-Capture ChIP-seq of native MNase-digested chromatin, with matching inputs collected. For EdU-seq, cells were released in S phase for 1.5h, followed by 30min EdU pulse and collection at 2h in S phase, followed by EdU IP. **B.** H3.3 and **D.** EdU at 2h in S enrichment profiles from WT (as reference) and HIRA KO rescue with YFP (control) and HIRA plasmid at blurred (n = 3274) and buried sites (n = 574), sorted by size and centered at their middle ±0.5Mb. **C.** H3.3 and **E.** EdU at 2h in S mean signal at blurred and buried sites between 60-160kb in length, centered in their middle ±0.5Mb, corresponding to the conditions described above. Enrichment relative to input was calculated at 10kb bins as z-score of log_2_ IP/input.

Next, we examined whether the recovery of the H3 variant pattern could restore firing at pre-existing replication IZs in early S phase. To test this, we monitored early replication in synchronized cells by following DNA synthesis using both EdU incorporation and PCNA staining (Figure 4A, as in (Gatto et al., 2022)). Upon rescue with HIRA, the proportion of cells in S phase 2h after G1/S release significantly increased, suggesting a recovery of early initiation (Supplementary Figure 4E). EdU-seq at 2h after release in S revealed early firing again restricted within blurred sites and significantly increased at buried sites (Figure 4D, E, Supplementary Figure 4F, G). Thus, our data are consistent with a restoration of early replication firing patterns concomitant with the re-establishment of H3.3/H3.1 balance.

### HIRA rescue recovers H3.3 pattern and early initiation at buried sites without compartment reversal

Finally, we performed Hi-C to assess if genome organization was also recovered upon HIRA rescue. We transfected HIRA KO cells with HIRA-YFP (HIRA) or YFP (control) plasmid (Supplementary Figure 5A) and validated the re-establishment of H3.3 targeting to blurred and buried sites by SNAP-capture ChIP-seq (Supplementary Figure 5B). Hi-C maps from control (YFP) and HIRA rescue showed very few compartment switches (Figure 5A, grey dots), echoed by the behaviour of both blurred and buried sites (Figure 5A, green and red dots, respectively). While buried sites switched more than expected by chance (7.7%, Figure 5B), only 5.3% switched from compartment B to A upon HIRA rescue (in contrast to 20% A to B upon HIRA KO). Furthermore, HIRA rescue recovered both H3.3 enrichment and early firing in buried sites remaining in B regardless of whether they had previously switched compartment from WT to HIRA KO (Figure 5C). Finally, following HIRA rescue, the restoration of H3.3 enrichment and early initiation in buried sites occurred to similar extents whether they switched from B to A or remained in B (Figure 5D, Supplementary Figure 5C, D). We thus conclude that recovery of early initiation zones at buried sites occurs irrespective of initial compartment, and when a switch had occurred there was no need to switch back.

**Figure 5.**
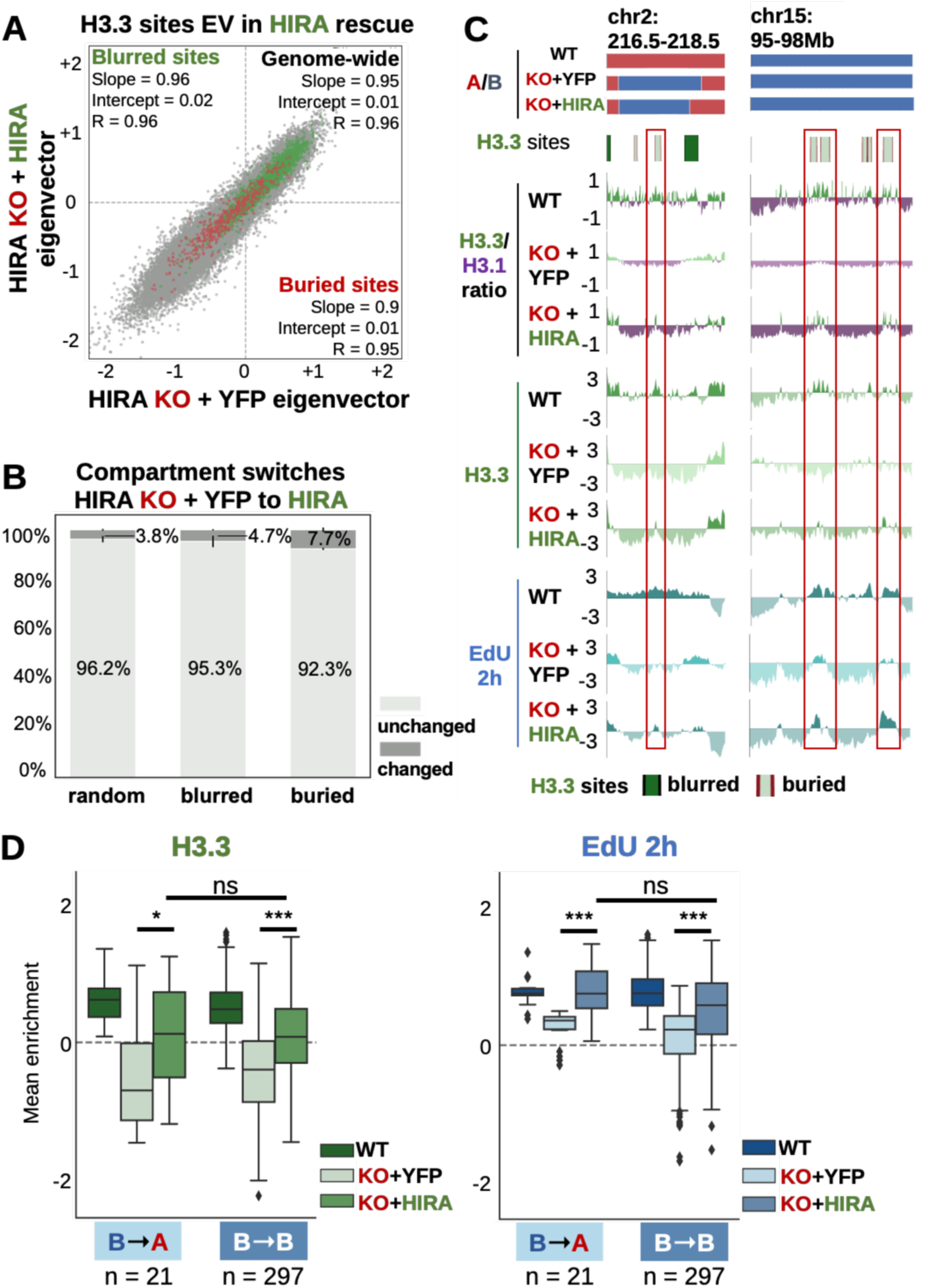
HIRA rescue recovers H3.3 pattern and early initiation at buried sites without compartment reversal. **A.** Comparison of EV1 distribution genome-wide (50kb bins, n = 40887, grey) and at blurred (n = 2382, green) and buried (n = 439, red) sites between HIRA KO + YFP (control) and HIRA rescue. Slope, intercept and R from linear regression are noted for each set. **B.** Proportions of blurred and buried sites which remain in the same compartment (unchanged) or undergo a switch (changed) from HIRA KO + YFP (control) to HIRA rescue. A set of randomised size-matched sites was quantified as control **C.** Compartment assignment, H3.3 site location (blurred/buried in dark/light green, respectively), H3.3/H3.1 ratio and enrichment of H3.3 and EdU 2h at representative regions from WT (as reference), HIRA KO rescue with YFP (control) and HIRA. Shown are a set of buried sites which shift from A-to-B after HIRA KO and remain B after rescue (left, cf. Supplementary Figure 3C, 5C) and a set of buried sites which are always in compartment B (right, cf. Figure 1B, 2A). Note that EdU 2h rescue is detected specifically at buried sites which increase H3.3 enrichment and H3.3/H3.1 ratio. **D.** Mean H3.3 and EdU 2h in S enrichment from WT (as reference) and HIRA KO rescue with YFP (control) and HIRA cells at buried sites that switched from B-to-A (n = 21) or remained in B (n = 297) from HIRA KO rescue with YFP (control) to HIRA. Enrichment relative to input was calculated at 10kb bins as z-score of log_2_ IP/input. H3.3/H3.1 ratio, H3.3 and EdU 2 in S enrichment (z-score of log_2_ IP/input ratio of cpm) are shown at 10kb bins smoothed over 3 non-zero bins. Two-tailed Mann-Whitney U test corrected for multiple testing by FDR (5% cut-off) was used to determine significance of differences between HIRA KO rescue with YFP (control) and HIRA or between HIRA KO + HIRA rescue signal in sites remaining in B or switching from B to A upon HIRA rescue. Significance was noted as: * (p<=0.05), ** (p<=0.01), *** (p<=0.001) for all comparisons.

## Discussion

In this work, we examined the link between local chromatin state, early initiation and higher-order genome folding (Figure 6). We distinguished early initiation zones based on their behaviour with respect to H3 variants following knock-out of the H3.3 chaperone HIRA and then examined how other features changed. First, in HIRA KO, we found that early initiation did not strictly follow changes in chromatin accessibility and histone PTMs. Second, by combining deletion and rescue experiments with Hi-C we revealed that HIRA defines early IZs irrespectively of their compartment organisation. We discuss implications for HIRA-dependent early IZ definition with respect to (i) the local chromatin environment and (ii) the recruitment of HIRA and (iii) compartment organisation in the context of regulating early replication.

**Figure 6.**
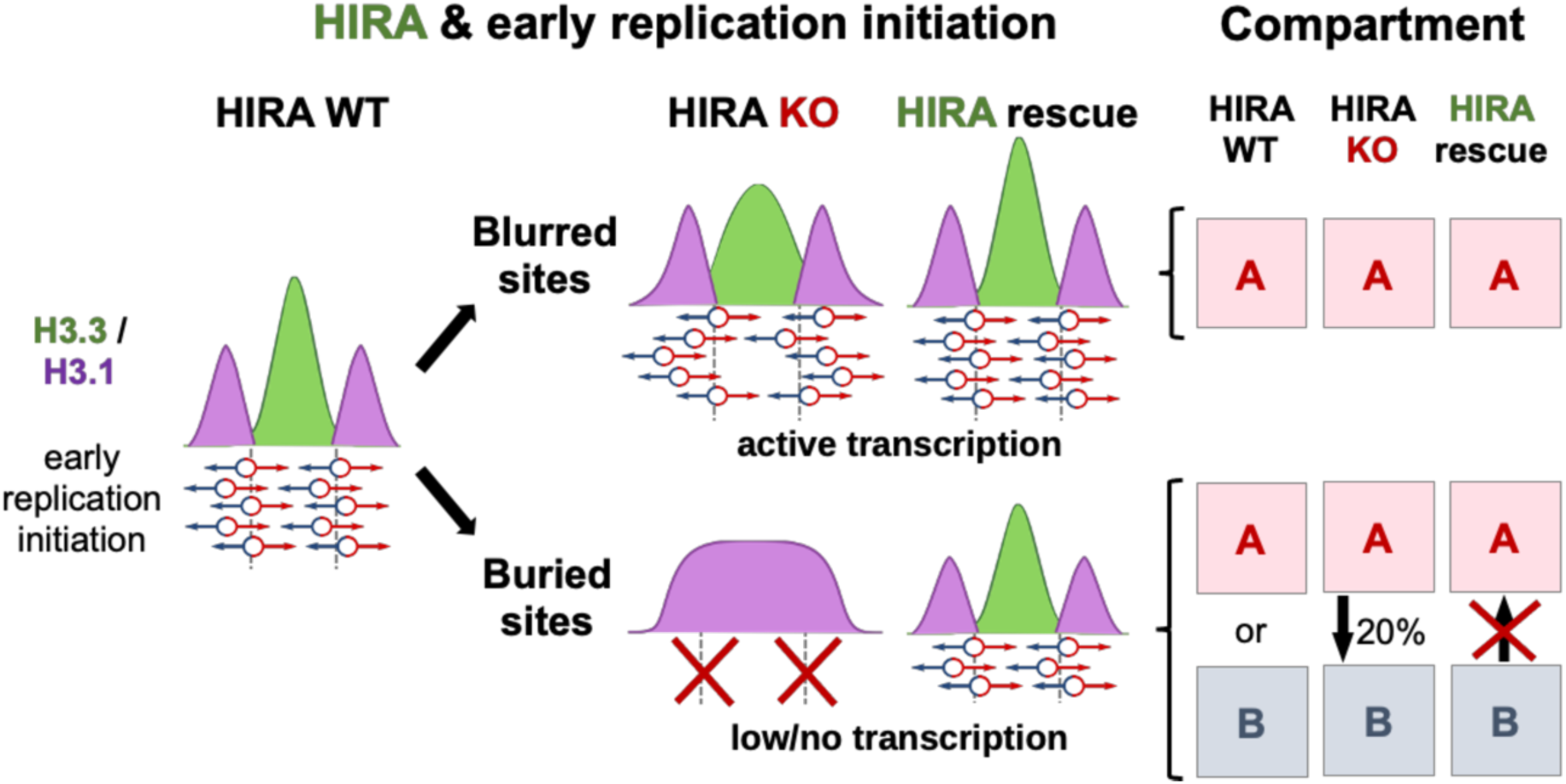
HIRA defines early initiation zones independently of their 3D organisation. HIRA-dependent early IZs are predominantly found in compartment A when actively transcribed (blurred sites). In contrast, early IZs can be found in both compartment A or B when they have low/no transcriptional activity (buried sites) in WT cells, indicating early initiation does not always occur in compartment A. Top: At blurred sites, H3.3 enrichment and early initiation become less precise in the absence of HIRA while remaining in compartment A. They regain sharpness upon rescue with HIRA also without changing compartment, indicating precision of initiation is not dependent on compartment identity. Bottom: Buried sites lose H3.3 enrichment and early firing in the absence of HIRA, but only a subset switches from compartment A to B. Furthermore, upon rescue with HIRA buried sites restore H3.3 enrichment and early firing without substantial switch from compartment B to A, indicating HIRA defined early replication initiation zones independently of their compartment organisation.

### HIRA defines early replication IZs independently of chromatin accessibility and histone marks

In line with other reports examining early initiation (Zhu and Kanemaki, 2024), we found that HIRA-dependent early IZs at the flanks of both blurred and buried sites show high accessibility in WT cells. In the absence of HIRA, accessibility increases within blurred and weakly decreases in buried sites. These changes do not match the loss of precision or detection of early initiation, respectively. In contrast, we reported increased accessibility at active gene bodies in a manner correlated with their transcriptional activity when HIRA was impaired (Karagyozova et al., 2024). Given that blurred sites are enriched in expressed genes (Gatto et al., 2022), this finding can explain their increased accessibility. Thus, these observations support a role for HIRA in defining early IZs independently of its role in regulating chromatin accessibility. Furthermore, our results also underline the fact that IZ accessibility on its own is not sufficient to ensure precision (at blurred sites) or timing (at buried sites) of early initiation.

In addition to accessibility, a defined set of histone marks have also been associated with early IZs (Hu and Stillman, 2023). Here, we found that both blurred and buried sites are enriched in active (H3K4me1/3, H3K27ac) and depleted in inactive (H3K9me3, H3K27me3) PTMs. Strikingly, these patterns remained similar in the absence of HIRA, unlike H3.3 and early initiation. Importantly, this finding parallels their preserved enrichment in compartment A/B in the absence of HIRA (Karagyozova et al., 2024). This demonstrates that HIRA does not contribute to early IZ definition by influencing the distribution of these marks. Interestingly, the histone methyltransferase MLL3/4 was reported to promote MCM2 enrichment at a set of H3K4me1 peaks. Notably, enrichment of H3.3 also occurs at these peaks which are lost together with early replication in the absence of MLL3/4 (Goekbuget et al., 2023). Thus, rather than the presence of the mark as key to define early initiation at these sites, we would like to propose to consider H3.3 deposition and its turnover. Indeed, the prediction of early replication based on active marks or accessibility, although highly correlated, is less robust than the use of H3.3 (Goekbuget et al., 2023; Halliwell et al., 2024). Here, we have demonstrated that the early initiation defect upon HIRA loss mirrored the impaired H3.3 distribution rather than accessibility or PTM patterns. In conclusion, our data put forward a key role for H3.3 deposition by HIRA in defining early IZ firing independently of the histone marks we assayed.

### HIRA rescue enables recovery of H3 variant enrichment defining sharp boundaries and early firing at IZs irrespective of compartment

The majority of HIRA-dependent early IZs are located at the boundaries of what we had previously coined blurred sites. We find these mainly in compartment A, where they remain in the absence of HIRA despite the reduced precision in both H3.3 enrichment and early replication (Gatto et al., 2022). At blurred sites, HIRA rescue reestablishes a sharp boundary of H3.3/H3.1 and restores early initiation profiles, again without changes in compartment. We propose that by interacting with RNA Pol II or via its ‘gap-filling’ mechanism (Ray-Gallet et al., 2011), HIRA recruitment could rely on the presence of active genes and increased accessibility in blurred sites. Thus, at blurred sites, while precision of early replication initiation is independent of compartment organisation, transcription could act as a memory to recruit HIRA back. In contrast, early initiation can also occur at the boundaries of buried sites which were coined this way because they completely disappeared in HIRA KO. They absolutely require HIRA for early initiation and in WT cells, they are predominantly in compartment B. Strikingly, HIRA rescue recovered both H3.3 enrichment and early initiation at buried sites despite their low/no transcriptional activity. Here, it is important to envisage how HIRA is recruited back given that we cannot invoke a transcription-linked marking as above. Compartment organisation is unlikely to support HIRA targeting to early IZs, given the disconnection we documented with H3.3 enrichment and early firing (discussed in more detail below). The fact that H3.3 enrichment can be precisely recovered within buried sites implies they must bear a clear demarcation which is (i) maintained ‘as a bookmark’ in the absence of HIRA and (ii) sufficient to guide it back following addback. On the one hand, given the capacity of HIRA to directly bind DNA (Kim et al., 2024; Ray-Gallet et al., 2011), it is interesting to consider that buried sites may have particular DNA properties which mediate this recruitment. Although replication initiation in metazoans does not occur at defined sites, sequence features like G-rich elements can promote early firing (Hu and Stillman, 2023). On the other hand, it is tempting to speculate that the ‘memory’ of buried sites may relate to PTM maintenance in the absence of HIRA. Additionally, buried site boundaries also remain accessible in the absence of HIRA. Apart from transcription, accessibility can be generated by chromatin remodellers, providing another potential mechanism for HIRA recruitment given its reported interaction with the SWI/SNF complex member BRG1 (Pchelintsev et al., 2013). Finally, the HIRA chaperone complex comprises three subunits: HIRA (Lamour et al., 1995), UBN1/2 (Banumathy et al., 2009; Tagami et al., 2004), CABIN1 (Balaji et al., 2009; Rai et al., 2011; Tagami et al., 2004). Although their stability requires the HIRA protein (Ray-Gallet et al., 2011), UBN1 and CABIN1 remain detectable in the nuclei of HIRA KO cells (Ray-Gallet et al., 2018). Since UBN1 can bind DNA (Ricketts et al., 2019), even small amounts retained locally could provide an alternative for ‘memory’ of the sites. Overall, our data demonstrates that HIRA/H3.3 deposition targeting is successfully restored upon rescue at both types of pre-existing H3.3 sites. While blurred sites could rely on RNA Pol II for this recovery, at buried sites this occurs independently of transcription. Importantly, we should emphasize that regardless of the type of IZ considered, H3.3 and early initiation recovery is independent of compartment organisation.

### HIRA-mediated H3.3 deposition at early IZs as a model to dissect the relationship between replication initiation control and 3D genome organisation

Disentangling the link between early/late replication and A/B compartments has proven challenging due to our limited understanding of the factors that govern them (Li et al., 2024; Vouzas and Gilbert, 2021). Here, we show that in the absence of HIRA, half of the buried sites previously in compartment A switch to B. This occurs specifically at non-transcribed sites without significant changes in expression. In contrast, buried sites in compartment A which were transcribed in WT cells remain there following HIRA deletion. Thus, loss of early initiation alone is not sufficient to result in a compartment change. Notably, the restoration in replication and H3.3 at buried sites did not require switching back from compartment B to A at our timescale. This implies that the role in HIRA in defining early IZs is independent of their compartment organisation. To our knowledge, comparison of compartment organisation between unperturbed and impaired early firing IZs has not been reported to date. However, a global impairment of temporal replication control occurs upon depletion of RIF1 (Cornacchia et al., 2012; Yamazaki et al., 2012) or MCM6 (Peycheva et al., 2022). Yet, only in the case of RIF1, reports of changes in compartments have been identified in a cell type-specific manner (Klein et al., 2021; Malzl et al., 2023). However, it remained unclear how switching corresponded to changes in early initiation. In contrast, DNA methylation can impact compartment organisation, but there is conflicting evidence whether this is accompanied by changes in RT (Du et al., 2021; Spracklin et al., 2022). We have recently demonstrated that HIRA plays a role in the local organisation of active genes and compartment A interactions (Karagyozova et al., 2024). Here, we show that this function is independent of its importance for early IZ definition. Thus, we also unveil distinct features in HIRA functions impinging independently on compartment A organization in a manner linked to transcription and the definition of early initiation zones.

In conclusion, our work demonstrates that HIRA defines early replication IZs independently of accessibility and histone H3 PTMs and irrespectively of the compartment they are in. We highlight how transcription-independent HIRA recruitment to early IZs provides a novel opportunity to understand how early replication and compartment organisation can be independently regulated.

## Materials and Methods

### Cell culture

We used HeLa cells stably expressing H3.1-SNAP-HA or H3.3-SNAP-HA which were knocked out for HIRA (KO, CRISPR/Cas9 HIRA KO) and transfected with YFP (control) or HIRA-YFP plasmids as in (Ray-Gallet et al., 2018). Cell lines were grown and synchronized at the G1/S transition using a double thymidine block as described in (Forest et al., 2024; Gatto et al., 2022), starting the first block 6h post-transfection (Figure 4A). All cell lines were tested negative for mycoplasma.

### Microscopy

To monitor replication, we performed 30min EdU pulse (as described below for EdU-seq) or PCNA staining in G1/S-arrested and 2 hours-released cells. We monitored transfection efficiency 48 hours after transfection by detecting YFP. Cells were fixed for 15min with 2% PFA. For PCNA labelling, we performed pre-extraction of soluble proteins with Triton-CSK for 5min, fixed in 2% PFA as above and then post-fixed cells with methanol for 20min at -20°C. Cells were blocked for with 5% BSA for 1h prior to immunofluorescence staining performed as in (Clément et al., 2018) with PCNA antibody (DAKO, M879, 1:1000). We detected EdU using the Click-iT EdU Cell Proliferation kit as described in (Gatto et al., 2022). Cells were co-stained with DAPI to label DNA. We imaged using a Zeiss Images Z1 fluorescence microscope with 63x and 40x oil objective lenses, ORCA-Flash4.0 LT camera (Hamamatsu) and MetaMorph software. We used ImageJ (Fiji) software for image visualization and analysis.

### SNAP capture-seq and EdU-Seq

We performed SNAP capture-seq of transfected G1/S-synchronised cells by double thymidine block as described in (Forest et al., 2024; Gatto et al., 2022). We carried out EdU labelling to map sites of ongoing synthesis from G1/S synchronized cells by a double thymidine block and released in S-phase as described in (Gatto et al., 2022). Ninety minutes after release of cells from the G1/S block, we performed a 30min pulse labelling by adding EdU (25μM) to the medium. We also performed SNAP capture-seq of transfected asynchronous cells (from Hi-C rescue, described below) as described in (Forest et al., 2024; Gatto et al., 2022) starting with 2 million cells per condition. We quantified and checked fragment size profile with Agilent 4200 TapeStation. We prepared sequencing libraries at the Next Generation Sequencing (NGS) platform from Institut Curie with the Illumina TruSeq ChIP kit and sequenced on Illumina NovaSeq 6000 (PE100).

### Hi-C

We performed Hi-C using the Arima Hi-C+ kit following the manufacturer’s instructions and as described in (Karagyozova et al., 2024), starting with three-five million asynchronous HeLa cells per condition (Supplementary Figure 5A). After ligation of Illumina TruSeq sequencing adaptors, we performed Arima QC2 (library quantification) to determine the number of amplification cycles for the library PCR using the KAPA Library Quantification Sample kit following the manufacturer’s instructions. Amplified libraries were sequenced as described above.

### Sequencing data processing

Sequencing data was processed from raw reads in FASTQ format as described in (Karagyozova et al., 2024). Analysis of the data was carried out by custom Python scripts also as described in (Karagyozova et al., 2024). For each sample, counts were read at 100bp or at 1kb resolution and aggregated to 10kb bins. Binned data (at 100bp, 1kb and 10kb) was normalized to the total number of mapped counts to generate cpm (counts per million). ChIP-seq data was normalized to matched input sample by computing a log_2_ ratio (IP/input), then scaled and centered by computing a z-score per chromosome. RNA-seq data was plotted as log_2_(cpm+1). Blurred (n = 3274) and buried (n = 574) site locations and H3.3+ OK-seq IZ (n = 5596) coordinates were obtained from (Gatto et al., 2022).

### Hi-C data processing

We processed Hi-C data as described in (Karagyozova et al., 2024). Matrices were masked using a custom set of blacklisted regions (Karagyozova et al., 2024) prior to normalisation. Matrices from the H3.1- and H3.3-SNAP cell lines were analysed independently and showed similar results. Coordinates and data from H3.1-SNAP cells were used for representative images where experiments were performed in both cell lines unless mentioned otherwise.

### Compartment analysis

Compartment analysis of Hi-C matrices was performed by eigenvector (EV) decomposition (Lieberman-Aiden et al., 2009) at 50kb resolution with cooltools (Open 2C et al., 2022) using GC content track to orient the sign of the first eigenvector (EV1). A/B compartment domains were defined for each sample as contiguous segments of the genome with the same EV1 sign. After masking the Hi-C data, we retained 2382 blurred and 439 buried sites that were used for compartment analysis. To assess their compartment identity, we re-binned the EV1 signal to 10kb and we calculated the mean EV1 value from each sample. To determine compartment switches of the sites, we compared their compartment assignment between WT and HIRA KO (WT-to-KO) or HIRA KO + YFP to HIRA KO + HIRA rescue (YFP-to-HIRA). The proportion of sites changing or remaining in the same compartment are represented by the mean and standard deviation of the two cell lines. A set of random sites matched for size distribution was generated for comparison. As an additional control for the different compartment distribution of the blurred and buried sites, we also generated two sets of sites matching the size and compartment distribution of blurred and buried sites. As a reference for genome-wide behaviour in the scatterplots, we used 50kb-binned EV1 values.

### Quantifications and statistical analysis

Statistical analysis was performed in pyhton using scipy (Virtanen et al., 2020). p-values were calculated by two-tailed Mann-Whitney U test. Multiple testing correction was performed by controlling the false discovery rate (FDR) using the Benjamini-Hochberg method. Differences with adjusted p-value <0.05 were considered statistically significant. Paired t-test was used to compare the differences in efficiency of transfection and S phase entry from cell counts.

### Materials

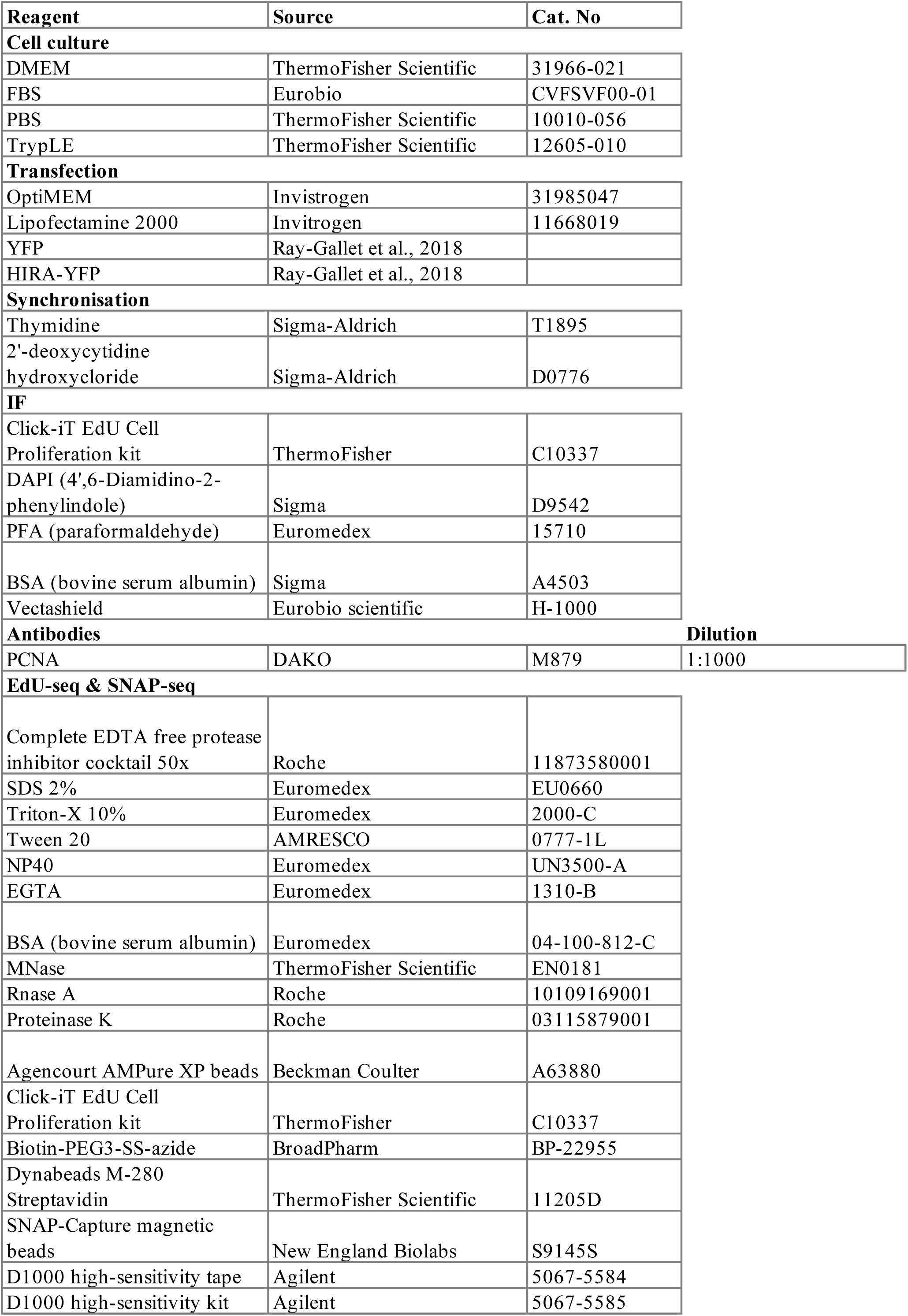
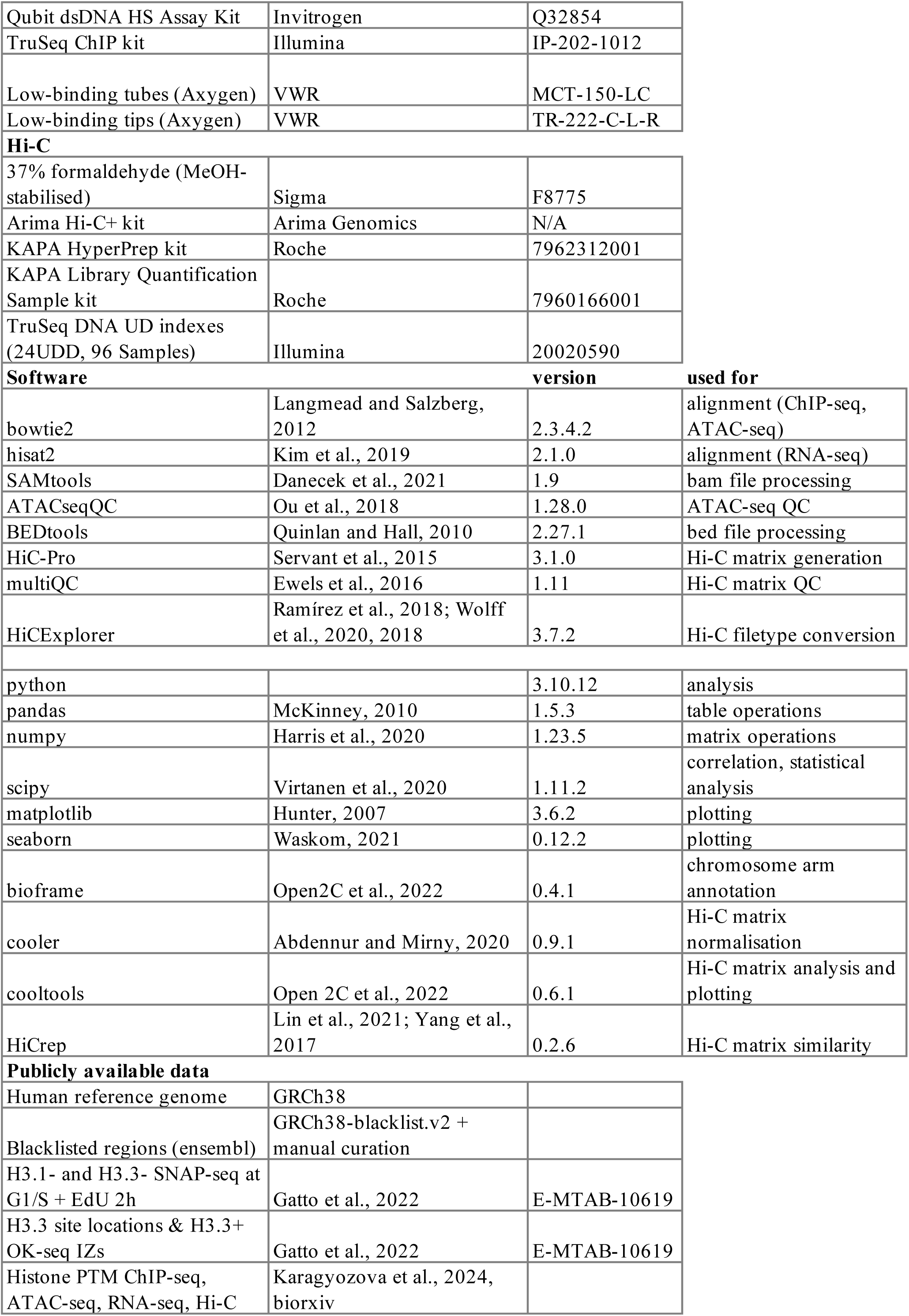

## Author contributions

G.A., J.P.Q., A.G. and T.K. conceived the overall strategy. T.K. and G.A. wrote the paper. T.K., A.F. and J.P.Q. performed the experiments. T.K. generated most of the figures and analysed data. G.A. supervised the work. M.A.M-R., A.G. and L.M. provided advice for the analysis. Critical reading and discussion of data involved all authors.

## Acknowledgements

We thank the members of UMR3664 and Almouzni team for helpful discussions. We thank Sébastien Lemaire for critical reading of the Methods and Dominique Ray-Gallet for constructive feedback on the model. We acknowledge the Cell and Tissue Imaging Platform PICT-IBiSA (member of France-Bioimaging ANR-10-INBS-04) of the UMR3664 and ICGex NGS platform of the Institut Curie.

This work was supported by the European Research Council (ERC-2015-ADG-694694 ‘‘ChromADICT’’), the Ligue Nationale contre le Cancer (Equipe labellisée Ligue), France and Agence Nationale de la Recherche, France (ANR-11-LABX-0044_DEEP, ANR-10-IDEX-0001-02 PSL, and ANR21-CE-11-0027 ‘‘CAFinDs’’). T.K. was supported by individual funding from H2020 MSCA-ITN – ChromDesign (Grant No. 813327) and La Ligue Nationale contre le Cancer (Grant No. TDLM23697). M.A.M-R. acknowledges support by the Spanish Ministerio de Ciencia e Innovación (PID2020-115696RB-I00 and PID2023-151484NB-I00).

---

**Supplementary Figure 1.**
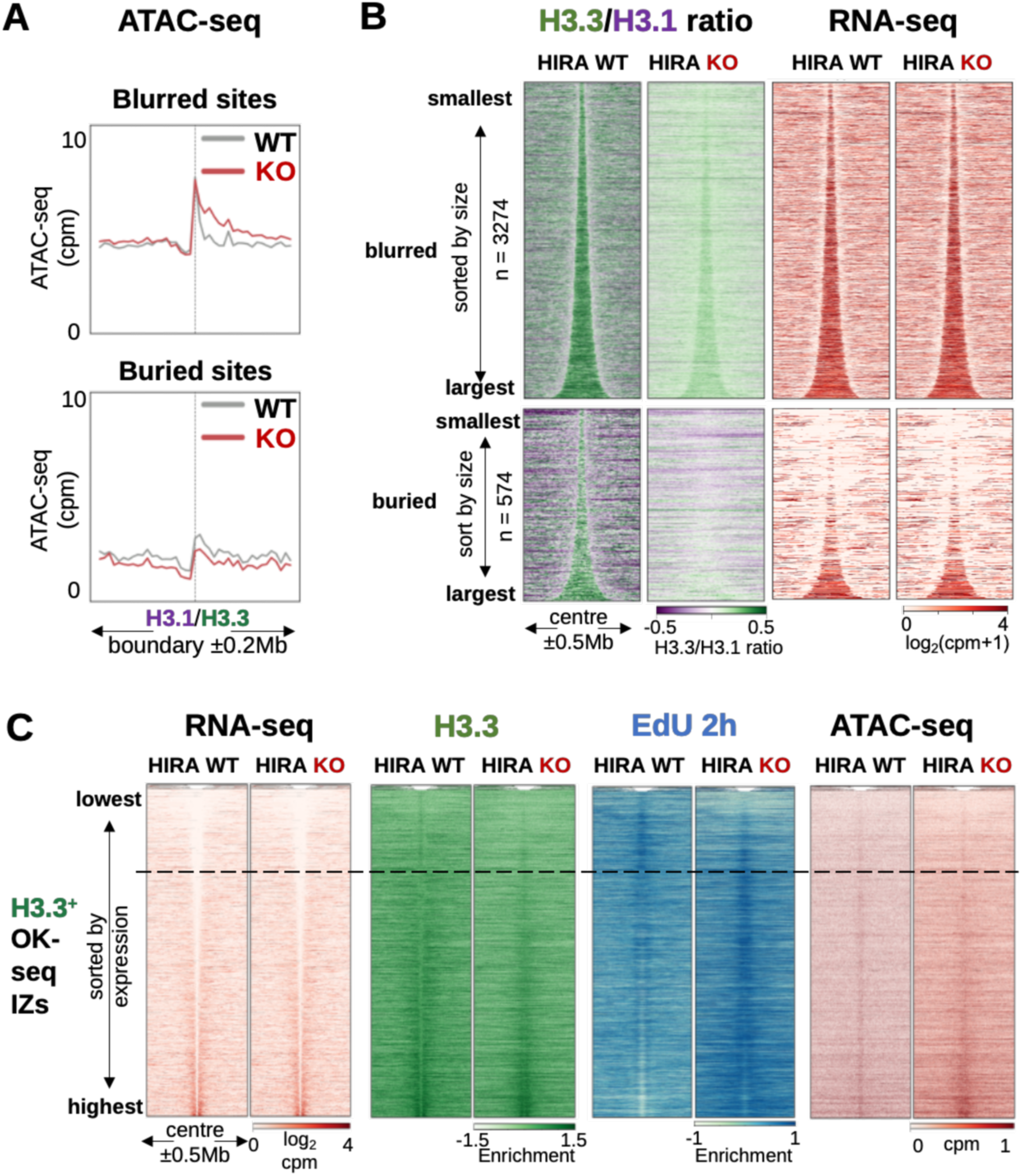
HIRA defines early replication initiation zones independently of its importance for chromatin accessibility. **A.** Mean ATAC-seq signal at 10kb bins from WT and HIRA KO cells centered at the H3.1/H3.3 boundaries of blurred (n = 3274) and buried (n = 574) sites ±0.2Mb. **B.** H3.3/H3.1 ratio and RNA-seq at 10kb bins from G1/S-arrested WT and HIRA KO cells at blurred and buried sites, sorted by size and centered at their middle ±0.5Mb. **C.** RNA-seq, enrichment of H3.3, EdU at 2h in S and ATAC-seq signal at 1kb bins from WT and HIRA KO cells at H3.3^+^ OK-seq IZs (n = 5596, as classified in (Gatto et al., 2022)), sorted by RNA-seq signal and centered at their middle ±0.5Mb. The dashed line represents bottom quartile of expression. H3.1, H3.3 and EdU 2h enrichment relative to input was calculated at 1kb bins as z-score of log_2_ IP/input. RNA-seq is plotted as log_2_(cpm+1) and ATAC-seq as cpm.

**Supplementary Figure 2.**
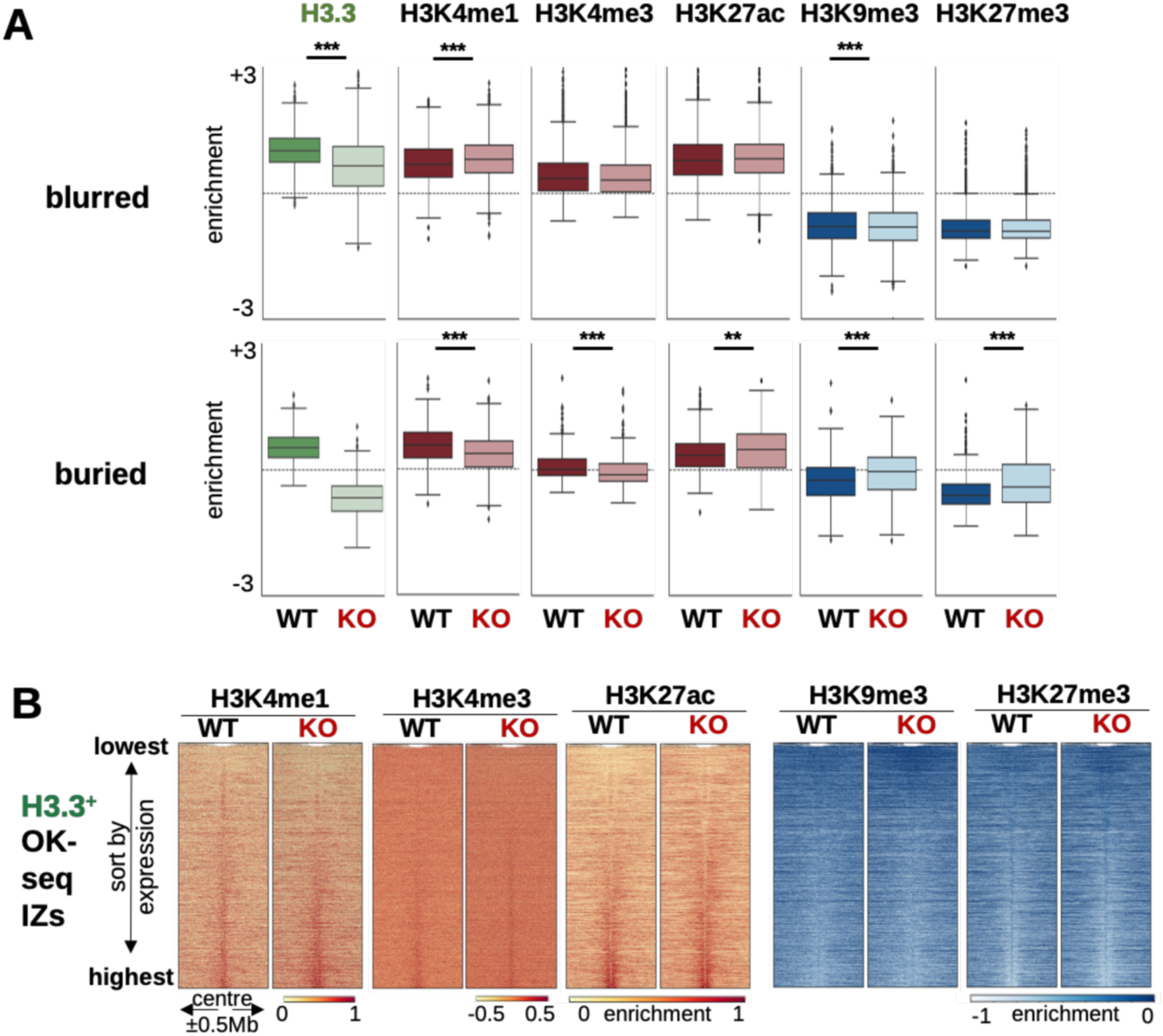
Impaired early IZ firing in the absence of HIRA is not associated with local H3 PTM redistribution. **A.** Mean enrichment of active (H3K4me1, H3K4me3, H3K27ac) and repressive (H3K9me3, H3K27me3) histone PTMs at 10kb bins from WT and HIRA KO cells at blurred (n = 3274, top) and buried (n = 574, bottom) sites. **B.** Active (H3K4me1, H3K4me3, H3K27ac) and repressive (H3K9me3, H3K27me3) histone PTM enrichment at 1kb bins from WT and HIRA KO cells at H3.3^+^ OK-seq IZs (n = 5596, as classified in (Gatto et al., 2022)), sorted by RNA-seq signal and centered at their middle ±0.5Mb. Enrichment relative to input was calculated as z-score of log_2_ IP/input. Two-tailed Mann-Whitney U test corrected for multiple testing by FDR (5% cut-off) was used to determine significance of differences between WT and HIRA KO. Significance was noted as: * (p<=0.05), ** (p<=0.01), *** (p<=0.001) for all comparisons.

**Supplementary Figure 3.**
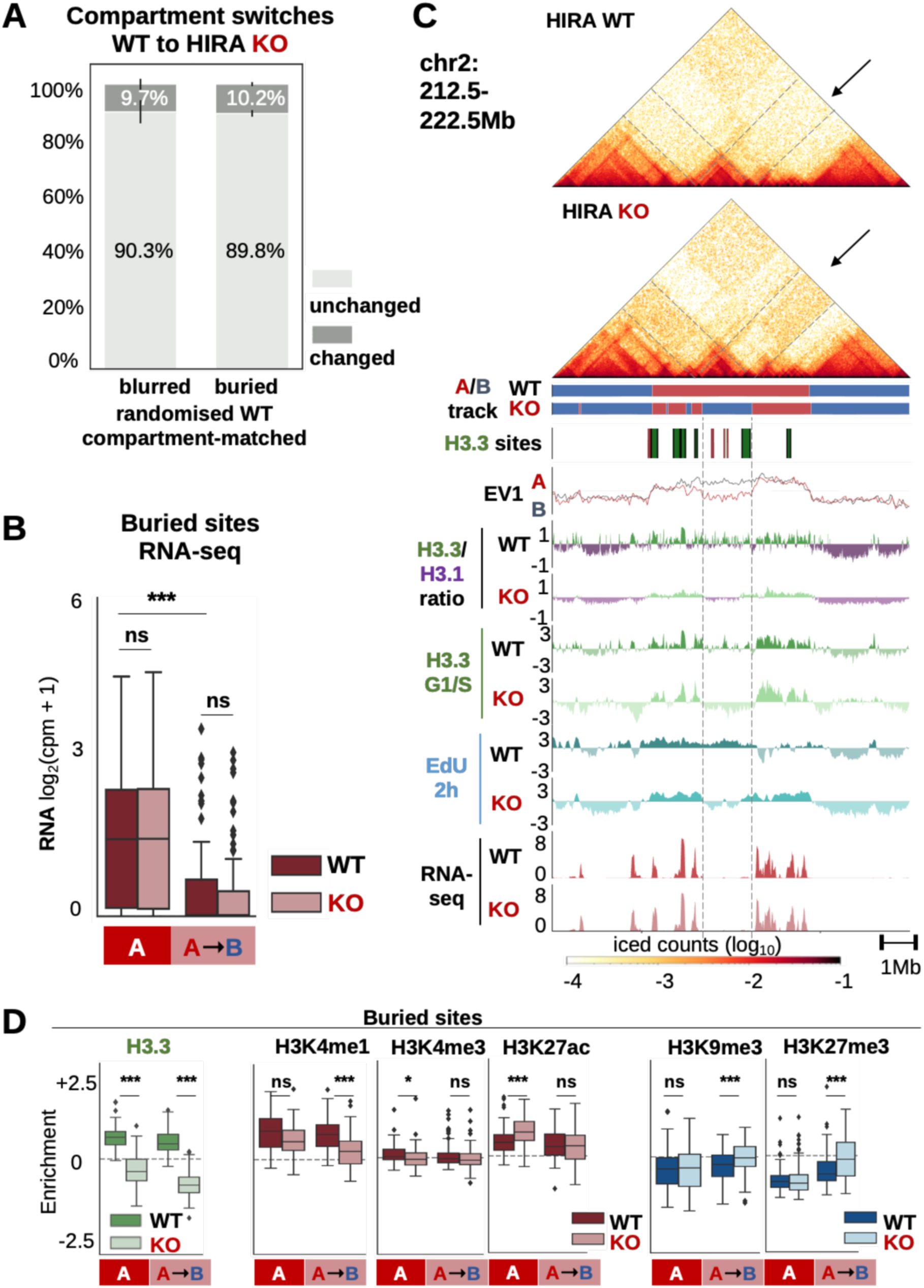
In the absence of HIRA, only non-transcribed early initiation zones switch from A to B compartment. **A.** Proportion of a size- and WT compartment assignment-matched set of blurred or buried sites which remain in the same compartment (unchanged) or undergo a switch (changed) from WT to HIRA KO cells. **B.** RNA-seq at buried sites which remain in compartment A (n = 92) or switch form A-to-B (n = 92) form WT to HIRA KO cells. **C.** Hi-C maps from WT and HIRA KO cells at a representative region switching compartment (chr2: 212.5-222.5Mb). Compartment track and blurred and buried site locations (dark and light green boxes, respectively) are noted below. EV1 signal (WT, grey, and HIRA KO, red), H3.3/H3.1 ratio, H3.3 enrichment, EdU at 2h in S and RNA-seq from WT and HIRA KO cells are shown at 10kb bins smoothed over 3 non-zero bins. Grey vertical lines and arrows denote a cluster of buried sites which are not transcribed and switch from compartment A to B concomitantly with losing EdU incorporation at 2h in S and H3.3 enrichment in HIRA KO. **D.** H3.3, active (H3K4me1, H3K4me3, H3K27ac) and repressive (H3K9me3, H3K27me3) histone PTM enrichment from WT and HIRA KO cells at buried sites which remain in compartment A or switch from A-to-B in HIRA KO cells. H3.3, H3 PTM and EdU enrichment are calculated as z-score of log_2_ IP/input ratio of cpm and RNA-seq is shown as log_2_(cpm+1), all at 10kb resolution. Two-tailed Mann-Whitney U test corrected for multiple testing by FDR (5% cut-off) was used to determine significance of differences between WT and KO. Significance was noted as: * (p<=0.05), ** (p<=0.01), *** (p<=0.001) for all comparisons.

**Supplementary Figure 4.**
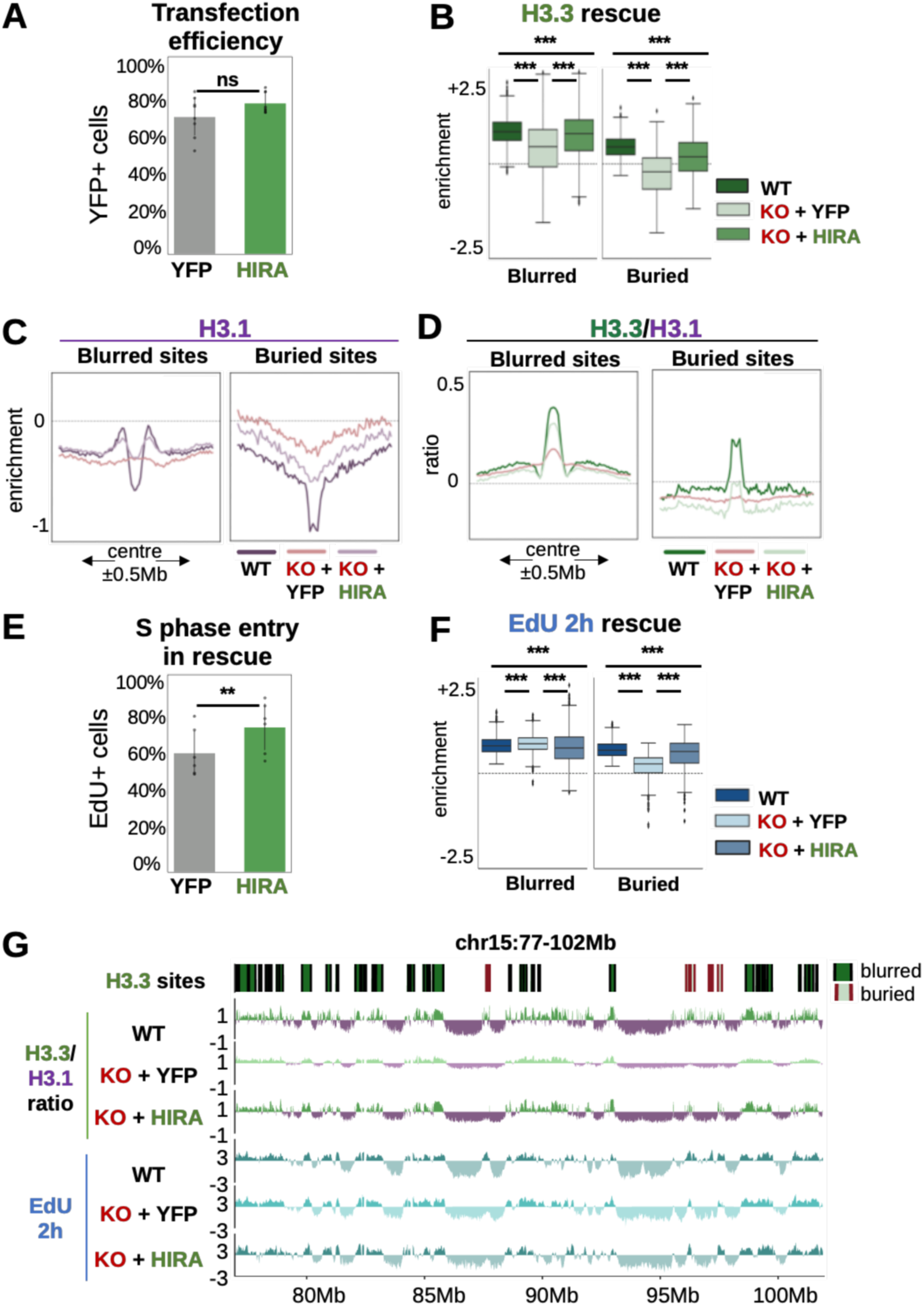
HIRA rescue reestablishes H3 variant pattern and early firing at blurred and buried sites. **A.** Proportion of YFP positive cells following transfection of HIRA KO cells with YFP (control) or HIRA-YFP. Mean, standard deviation and values of 8 independent experiments are shown. **B.** H3.3 (left) and enrichment from WT (as reference) and HIRA KO cells rescued with YFP (control) or HIRA at blurred (n = 3274) and buried (n = 574) sites. **C.** H3.1 enrichment from WT (as reference), and HIRA KO cells rescued with YFP (control) and HIRA at blurred and buried sites between 60-160kb in length, centred in their middle ±0.5Mb. **D.** H3.3/H3.1 ratio plotted as described for H3.1 enrichment above. **E.** Proportion of cells in early S phase 2h post-release from G1/S arrest from HIRA KO cells transfected with YFP (control) or HIRA-YFP. Mean, standard deviation and values of 6 independent experiments are shown. **F.** EdU at 2h in S (right) enrichment from WT (as reference) and HIRA KO cells rescued with YFP (control) or HIRA at blurred and buried sites. **G.** H3.3/H3.1 ratio and EdU at 2h in S from WT (as reference) and HIRA KO cells rescued with YFP (control) and HIRA along a representative region (chr15:77-102Mb) containing blurred and buried sites (denoted in dark and light green on top of the tracks, respectively). H3.3 and EdU enrichment were calculated at 10kb bins as z-score of log_2_ IP/input. Two-tailed Mann-Whitney U test corrected for multiple testing by FDR (5% cut-off) was used to determine significance of differences between WT, HIRA KO + YFP (control) and HIRA KO + HIRA rescue. Paired t-test was used to determine significance of differences between proportion of YFP+ or EdU+ cells following HIRA or YFP transfection. Significance was noted as: * (p<=0.05), ** (p<=0.01), *** (p<=0.001) for all comparisons.

**Supplementary Figure 5.**
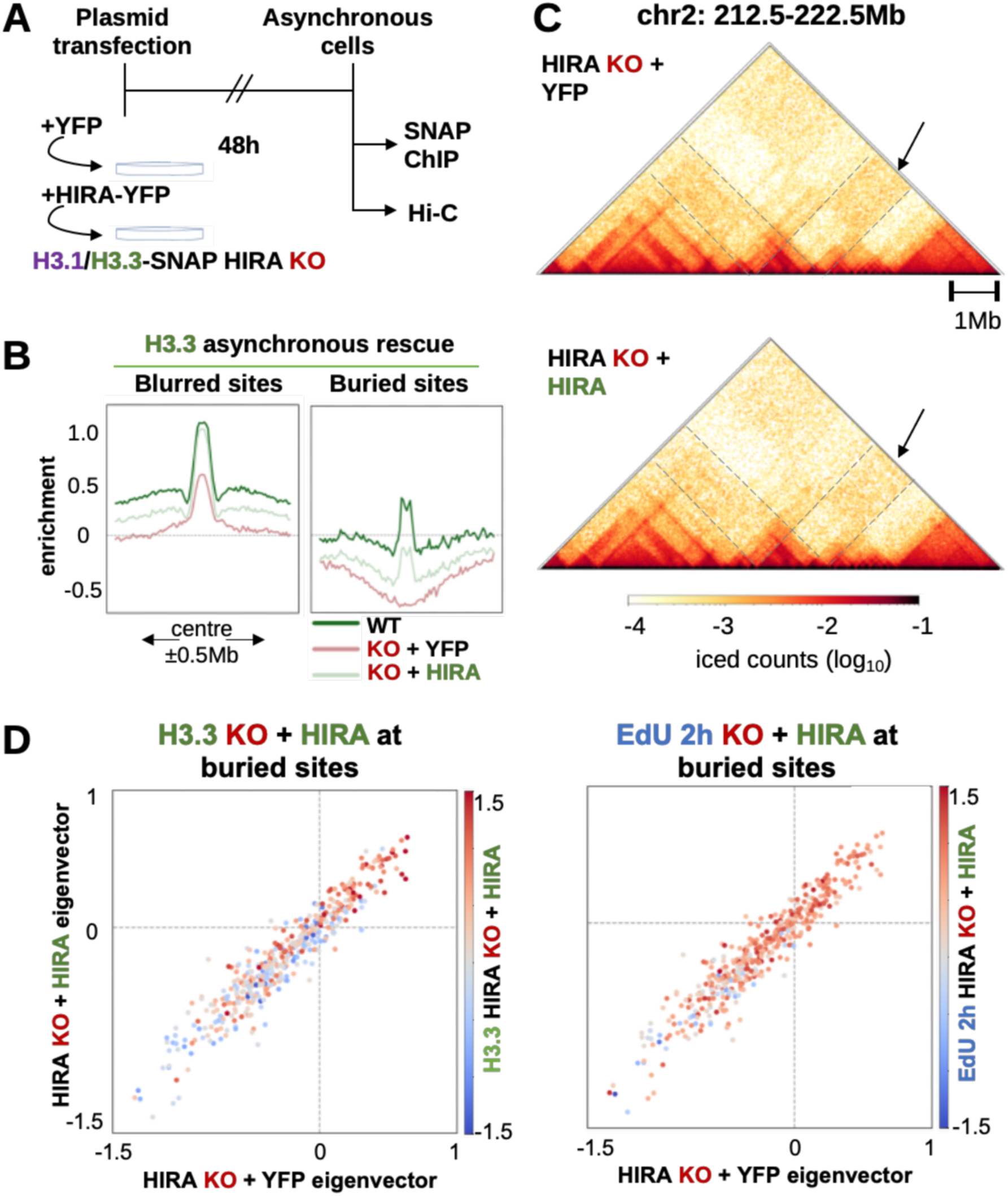
HIRA rescue recovers H3.3 enrichment and early firing at buried sites without compartment reversal. **A.** Scheme of experimental strategy to perform HIRA rescue in asynchronous cells to assay total H3.1/H3.3-SNAP and 3D genome organisation. Asynchronous cells constitutively expressing H3.1- or H3.3-SNAP were transfected with YFP (control) or HIRA-YFP plasmid for 48h. Total H3.1- and H3.3-SNAP were assayed by SNAP-Capture ChIP-seq of native MNase-digested chromatin, with matching inputs collected. Compartment organisation was assayed by Hi-C. **B.** Mean H3.3 enrichment from asynchronous WT (as reference) and HIRA KO cells rescued with YFP (control) and HIRA at blurred (n = 3274) and buried (n = 574) sites from 60-160kb in length. **C.** Hi-C maps from HIRA KO cells rescued with YFP or HIRA of a representative (chr2: 212.5-222.5Mb, cf. Figure 5C, Supplementary Figure 3C). Grey vertical lines and arrows denote a cluster of buried sites which switch from compartment A to B upon HIRA KO but do not revert from B to A after HIRA rescue despite increasing H3.3 and EdU 2h enrichment (Figure 5C). **D.** Scatterplots of mean EV1 value at buried sites (n = 439) from HIRA KO + YFP (control) and HIRA rescue coloured by their mean enrichment of H3.3 (left) or EdU at 2h in S (right) in HIRA KO + HIRA rescue. H3.3 and EdU 2h in S enrichment was calculated at 10kb bins as z-score of log_2_ IP/input.

